# *PIP2;1* aquaporin promotes early stomatal closure in grapevine leaves during water stress

**DOI:** 10.64898/2026.01.29.702672

**Authors:** Caetano Albuquerque, Mina Momayyezi, Cecilia B. Aguero, Celeste Arancibia, Ryan Stanfield, Mily Ron, Andrew Walker, Megan K. Bartlett, Christine Scoffoni, Andrew J. McElrone

**Author notes:** Authors contributed equally (Caetano Albuquerque, Mina Momayyezi).

## Abstract

Aquaporins play a key role in plant responses to drought. Our previous work showed limited embolism in grapevine leaves under mild water stress and suggested that the outside-xylem water pathway plays a dominant role in reducing leaf hydraulic conductance (*K*_leaf_) during dehydration. We used CRISPR-Cas9 to knockout the *PIP2;1* aquaporin encoding gene in *Vitis vinifera* cv. Chardonnay to study how leaf function during dehydration is affected by this aquaporin isoform. We measured functional responses like stomatal and photosynthetic responses as well as *K*_leaf_ to compare wild-type and two independent *PIP2;1* knockout lines. Under moderate drought, mutants maintained greater stomatal conductance (*g*_s_) and photosynthetic rates as Ψ_w_ declined. No significant differences were observed in mesophyll conductance (*g*_m_) across genotypes, however, mutants exhibited slightly higher values under moderate drought. Interestingly, all lines exhibited similar *K*_leaf_ vulnerabilities to drought. Our findings show that *PIP2;1* induces earlier stomatal closure during dehydration while not modulating *K*_leaf_ responses across genotypes. This rapid response in WT plants would prevent further water loss that would lead to higher xylem tensions that can lead to embolism. These findings show that multiple mechanisms collectively limit leaf gas exchange and water loss during dehydration, enhancing our understanding of plant resilience to changing environments.

## INTRODUCTION

Plants rely on an adequate water supply to leaves to maintain gas exchange and carbon capture for growth and development. Grapevines are an economically important crop worldwide and a key model system for plant hydraulics, largely cultivated in regions characterized by hot, dry summers where regulated water deficits are commonly applied to improve wine grape quality. Under these conditions, understanding the drivers of leaf water transport and stomatal conductance during drought and subsequent recovery is critical for both whole-plant physiology and crop management. Water vapor diffuses as a gas through stomata in exchange for atmospheric CO_2_ used in photosynthesis; as soil water becomes less available and/or atmospheric conditions become drier, stomata progressively close to limit further water loss at the expense of reduced photosynthesis. The challenge of balancing leaf hydraulic supply with atmospheric demand is intensified by the highly dynamic environment to which leaves are exposed, including increasing vapor pressure deficit (VPD), heat waves and soil drying, all of which constrain gas exchange and photosynthesis and can ultimately compromise yield (Dai, 2013; Grossiord et al., 2020). The regulation of stomatal conductance during dehydration is therefore central to preventing excessive xylem tensions that could otherwise lead to irreversible embolism (Scoffoni et al., 2023). In grapevines, loss of stomatal conductance (*g*_s_) and leaf hydraulic conductance (*K*_leaf_) precedes embolism inception in leaves (Albuquerque et al., 2020). These responses occur in a range of mild to moderate dehydration that plants commonly experience in the field (Williams & Araujo, 2002; Gambetta et al. 2020). Identifying the mechanisms that control these early hydraulic and stomatal responses, including the role of aquaporins, is thus essential for developing grapevines that are more productive and resilient in rapidly changing environmental conditions and during extreme weather events.

As water moves in the leaf, it travels first through the xylem in petioles and lamina veins (*K*_x_) and then reaches the outside-xylem pathways (*K*_ox_) made of living cells of the bundle-sheath and mesophyll. Thus, *K*_leaf_ is composed of both pathways: 1/*K*_leaf_ = 1/*K*_x_ + 1/*K*_ox_. Decreases in *K*_x_ under drought stress were assumed to be driven primarily by xylem embolism formation, but much work over the last decade has found leaf xylem is relatively resistant to embolism formation and that the living cells in the outside-xylem pathways are the main drivers of *K*_leaf_ decline during dehydration (Trifiló et al., 2016; Scoffoni et al., 2017, 2018; Albuquerque et al., 2020; Corso et al., 2020). Further, efforts using spatially-explicit models of outside-xylem leaf water transport points towards changes in cell membrane permeability associated with aquaporin activity as responsible for *K*_leaf_ decline during dehydration (Buckley, 2015; Buckley et al., 2015a).

Aquaporins (AQP) are protein channels embedded in plant plasma membranes and facilitate water movement between plant tissues (Chaumont and Tyerman, 2014). The plasma membrane intrinsic proteins (PIPs) are highly conserved and classified into two main groups, *PIP1*s and *PIP2*s (Bienert et al., 2018). Transport functionality of *PIP1*s and *PIP2*s has been characterized by heterologous expression in *Xenopus laevis* oocytes and *Saccharomyces cerevisiae*; by assessing both water transport activity and subcellular localization showed that only *PIP2*s mediate substantial transmembrane water fluxes (Yaneff et al., 2015). It has also been shown that *PIP1*s and *PIP2*s can form hetero-oligomers and that some *PIP1*s require a *PIP2* partner to reach the plasma membrane, suggesting the role of these interactions in *PIP* trafficking and function (Yaneff et al. 2015, Jozefkowicz et al. 2017, Bienert et al. 2018).

In grapevines, the *PIP* subfamily consists of eight members divided between *VvPIP1*s and *VvPIP2*s, which have been associated with water relations of various organs of grapevines under different conditions. In leaves of *Vitis vinifera* cv. Chardonnay, experiments using mercury chloride to inhibit aquaporin expression showed that the expression of *VvPIP 2;1* was correlated with declines in *K*_leaf_ and *g*_s_ during early dehydration and recovery (Pou et al., 2013). In leaves of Richter-110 (*V. berlandieri x rupestris*), aquaporin expression of seven isoforms including *VvPIP 2;1* changed depending on the level of water stress; they were down-regulated under moderate and up-regulated under severe water stress (Galmés et al., 2007). Daily fluctuations in the expression of aquaporin isoforms including *VvPIP2;1* were observed in leaf laminas and petioles of Chardonnay, decreasing during the day in conjunction with leaf water potentials (Shelden et al. 2017). Greater expression of *VvPIP1*s and *VvPIP2*s in roots of grapevine rootstocks was associated with higher vigor of the scion, and greater aquaporin expression in fine roots was associated with their increased root hydraulic conductivity compared to mature, more suberized root zones (Gambetta et al., 2012; 2013). The same aquaporins were shown to be coordinated with root hydraulic conductivity in two grapevine varieties with contrasting responses to water stress (Vandeleur et al., 2009). In grapevine roots, overexpression of VvPIP2;4N and VvPIP2;2 altered root water uptake and hydraulic conductivity primarily under well-watered conditions (Perrone et al., 2012). Overexpression of a tonoplast aquaporin (*SlTIP2;2*) modulated whole-plant responses in tomato, resulting in increased transpiration, growth and fruit yield under water and salt stress (Sade et al., 2009). These studies implicate aquaporins as key regulators of water transport and stress responses in grapevine and other crops, underscoring the need for approaches that resolve specific aquaporin isoforms as targets for improving crop performance and resilience under rapidly changing environmental conditions.

Several techniques have been used to measure and manipulate aquaporin expression in grapevines, including the use of mercury chloride to inhibit aquaporin activity. However, mercury chloride can induce cell toxicity and does not target specific aquaporin to be knocked out (Sabir et al., 2021). In grapevine roots, overexpression of *VvPIP2;4N and VvPIP2;2* did not alter root water uptake and hydraulic conductivity under water stress, but only under well-watered conditions (Perrone at al., 2012). Thus, despite extensive work on aquaporins in grapevine, the isoform-specific role of VvPIP2;1 in regulating leaf hydraulics and gas exchange during dehydration remains to be resolved. Newer molecular technologies like CRISPR-Cas9 allow targeting specific aquaporin isoforms with gene-editing to manipulate them and study their functional role of grapevine responses to stress. Here, we used CRISPR-Cas9 to knock out the expression of a *VvPIP2;1* isoform and evaluated its role in leaf responses to drought stress in terms of gas exchange, photosynthetic response and leaf hydraulics in *Vitis vinifera* cv.

Chardonnay, one of the most globally cultivated wine grapes. We hypothesized that: i) CRISPR-Cas9 would effectively knockout *VvPIP2;1* and impact leaf functional responses to drought; ii); *VvPIP 2;1* mutants would exhibit different responses to progressive dehydration as *K*_leaf_ and stomatal conductance declines are regulated in the outside-xylem pathways; iii) *VvPIP 2;1* mutants would have altered responses in other leaf functions like photosynthesis and water potential.

## MATERIAL AND METHODS

### 1. Plant material

#### 1.1. DNA editing, guide RNA design and CRISPR-Cas9 plasmid construction

CRISPR guide RNAs were designed to target the first and second exon of aquaporin *PIP2;1* VIT_13s0019g04280 in PN12x (https://plants.ensembl.org/Vitis_vinifera/Info/Index). In order to clone two gRNA expressing cassettes and minimize repetition, we modified pMR217 (Ritter et al. 2017) and replaced the AtU6-26 promoter with a synthetic promoter based on the consensus sequence of the 3 U6 promoter variants present in the *Arabidopsis* genome (Waibel et al. 1990). To create this vector, we amplified pMR217 with primers U6_Chimera for (ATTGGGGTCTTCGAGAAGACCTGTTTTAG) and U6_Chimera.Rev2 (GATTGTTTTCAACTTGAAGAAGAAAAAAAGAGCCTGCTTTTTTGTAC) in reverse orientation to remove the existing AtU6-26 promoter. Then to generate the U6 synthetic promoter we used 2 oligos: U6p_Oligo.For (AAGTTGAAAACAATCTTCAAAAGTCCCACATCGATCAGGTGATTATAGCAGCTTAGTTT) and U6p_Oligo.Rev (CTCGAAGACCCCAATCACTACTTCGACTCTATCATTATATAAACTAAGCTGCTATA TATC), that were annealed and extended using Phusion Taq (New England Biolabs). The resulted double stranded DNA fragment was cloned into the pMR217ΔAtU6-26 promoter using the in-fusion reaction (Takara) to generate pMR299.

We used the CRISPOR software (Concordet et al. 2018) ranked by efficiency score according to Moreno-Mateos et al. (2015), to choose 2 gRNAs targeting the *VvPIP2-1* gene. The cloning of the guides into pMR299 and pMR218 (Ritter et al. 2017) was performed according to Fauser et al., 2014. Two 23 bp overlapping oligos, representing the guide plus 4bp overhang were annealed and cloned into *Bpil*-digested pMR299 or pMR218 to generate pMR472 and pMR473 respectively. The resulting plasmids were used in an LR recombination (Thermo Fisher Scientific, Waltham, MA, USA) to transfer the U6Cons-PIP2-1_G1 and AtU6-26p-PIP2-1_G2 sgRNAs cassette into pMR290 binary vector (Bari et al., 2019) containing 2x35SΩ-Cas9 to generate pMR474 (Figure 1). This vector was subsequently introduced into *A. tumefaciens* strain EHA105 for *in vitro* transformation of embryogenic cells of Chardonnay clone 4, as in Agüero *et al*. (2006).

**Figure 1.**
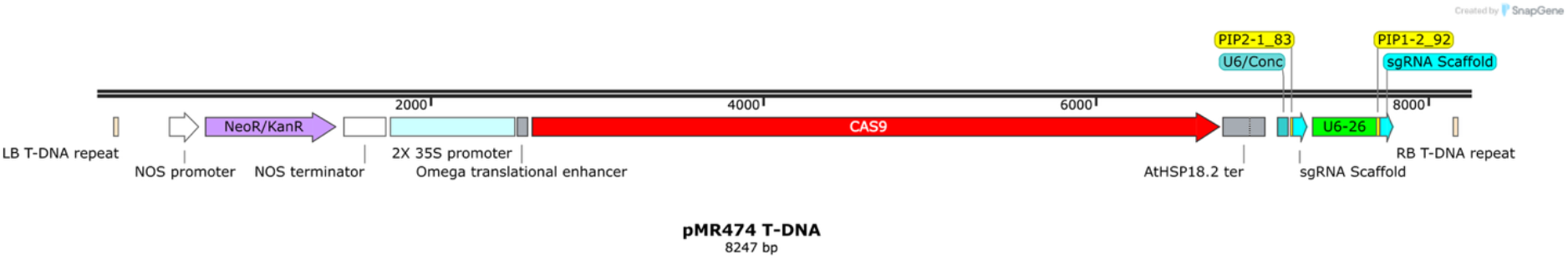
Schematic representation of the T-DNA in pMR474.

#### 1.2. Sequence analysis and plant selection

Eleven independent lines obtained from the *in vitro* transformation were acclimated to greenhouse conditions along with 2 wild type (WT) lines that developed from the same embryogenic callus (*i*.*e*. treated but not transformed) (see details in Table 1). Based on preliminary gas exchange measurements, two mutant lines (9 and 15) and one WT were selected from the original set for in-depth physiological and gene expression analysis. Both mutant lines were selected for showing the greatest differences from WT Chardonnay. Genomic DNA from these lines was extracted from leaves using the method described by Riaz *et al*. (2008). PCR products obtained using primers flanking the target sites were Sanger sequenced by QuintaraBio (http://www.quintarabio.com/services). The PCR conditions were 95 °C for 10 min, followed by 35 cycles of 95 °C for 45 sec, 60 °C for 30 sec, 72 °C for 90 sec, and a final step of 72 °C for 10 min using Applied Biosystems AmpliTaq Gold. Insertions and deletions were identified using the TIDE algorithm (Brinkman *et al*. 2014). The sequencing of genomic DNA of the independent lines showed that Crispr-Cas9 induced deletions at the first targeted site in both alleles while inducing insertions in the second site, but with lower efficiency. Line 9 and 15 have mutations in the two targeted sites, but they are not the same mutations (Table 1).. Selected lines (WT, 9 and 15) were clonally propagated from green cuttings, which were then rooted and planted in 1 L pots filled with a 1:1:1 Yolo sandy loam soil/perlite/peat mix. After a month, the plants were transplanted to 4 L pots and cut back to two buds to ensure the uniform growth of two shoots per plant. Grapevines were grown in a greenhouse at 18-25 °C and an 18-hour photoperiod with supplemental lighting (PPFD = 500 µmol m^-2^ s^-1^).

**Table 1.**
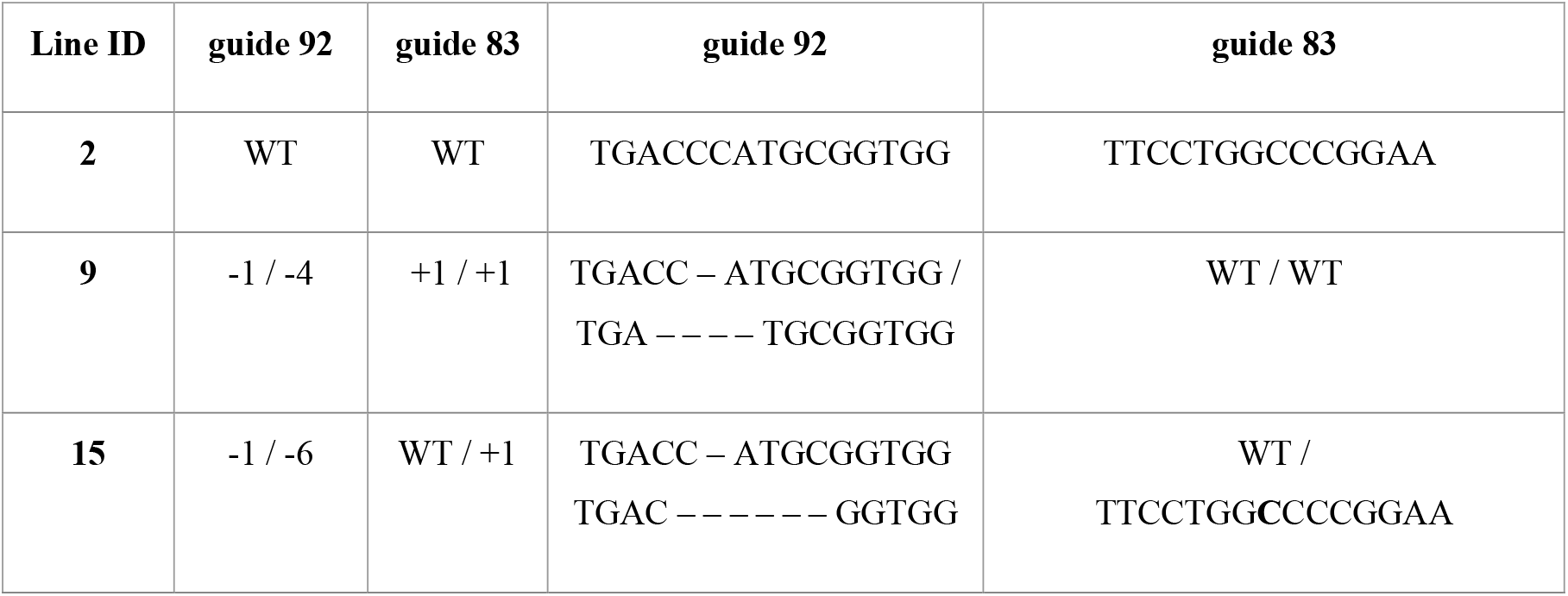
Indels elicited by an individual Cas9:gRNA92 and 83 in transgenic lines, sequence in WT and modifications in lines 9 and 15 are shown in the right columns.

#### 1.3. Drought treatments

A total 45 plants with 15 replications per line (WT, 9, 15) were kept well-watered for four weeks. Subsequently, the 15 replications per line were randomly assigned to one of three irrigation treatments: well-watered, mild drought and moderate drought using a completely randomized block design with five blocks. Each block contained one replicate of each irrigation treatment per plant line. To achieve the target water status, midday leaf water potentials (Ψ_leaf-midday_) were monitored daily. This was measured using a pressure chamber (PMS Instrument Company, Model 1505D) on the leaf immediately below the one used for the gas exchange measurement, between 10 AM to 12 PM. The predawn leaf water potential (Ψ_leaf-predawn_) was also measured on two leaves below the leaf measured for gas exchange, before sunrise, to assess drought stress and the effectiveness of the irrigation treatments. Plants were subjected to a rapid dry-down protocol by withholding water until they reached Ψ_leaf-midday_ targets of well-watered plants were maintained at a leaf water potential Ψ_leaf-midday_ > –1 MPa; mild at –1 MPa< Ψ_leaf-midday_ < –1.2 MPa; and moderate at –1.2 MPa< Ψ_leaf-midday_ < –1.4 MPa, reflecting typical drought conditions in commercial vineyards. These target leaf water potential ranges were established based on the methodology described by Albuquerque et al. (2020). To maintain the target water potential for each treatment, a subset of pots was monitored by weight, with water added in mL to compensate for evapotranspirational water loss. On average, well-watered plants required daily irrigation, mild stress plants were watered every 2–3 days, and moderate stress plants were watered only once per week. After one week of maintaining the target water status for each irrigation treatment, instantaneous leaf gas exchange and photosynthetic curve measurements were performed. Furthermore, aquaporin expression was analyzed under well-watered and moderate drought conditions, and *K*_leaf_ analysis was conducted using well-watered plants. Details of these procedures are provided in the subsequent sections.

### 2. Measurements

#### 2.1. Aquaporin expression

Leaves with petioles adjacent to the ones used for gas exchange measurements were collected at midday (1-2 PM) and the following morning (6 AM-6:30 AM). Total RNA was extracted using the CTAB method (Blanco-Ulate et al. 2013). The RNA pellet was further purified using the Quick-RNA Miniprep Kit (Zymo Research). cDNA was synthesized from the prepared RNA using M-MLV Reverse Transcriptase (Promega).

qRT-PCR was performed on a StepOnePlus PCR System using Fast SYBR Green Master Mix (Applied Biosystems) and the following primers:

*VvPIP1;1* F - TGGTGCGGGTGTAGTGAAGG R - AGACAGTGTAGACAAGGACGAAGG *VvPIP2;1* F - CAGGAGCACCACTCATGTATG R - TCATGCCCTCATACATATCAATAAC *VvPIP2;2* F - AAAGTTTGGGACGACCAGTG R - TTTTTAGTTGGTGGGGTTGC *VvPIP2;3* F - GCCATTGCAGCATTCTATCA R - TCCTACAGGGCCACAAATTC *VvTIP1;1* F - CATTGCCGCCATCATCTAC R - AGAAATCTCAACCCCACCAG *VvTIP2;1* F - GGAGGAAGAGCAAGTTGTGC R – GCACATCACCAACCTCATTC Actin: F - CTATGAGCTGCCTGATGGGC R – GCAGCTTCCATCCCAATGAG (Shelden et al. 2017, Jones et al. 2014). All qRT-PCR reactions were performed as follows: 50°C for 2 min, 95°C for 10 min, followed by 40 cycles of 95°C for 15 s and 60°C for 60 s. The 2-ΔΔCT method (Livak and Schmittgen 2001) was applied to analyze the expression levels of the genes.

#### 2.2. Leaf gas exchange

##### 2.2.1. Photosynthetic measurements

The instantaneous net assimilation rate (*A*_n_), stomatal conductance (*g*_s_) and the intercellular airspace CO_2_ concentration (*C*_i_) were measured on the 7^th^ or 8^th^ leaf per vine using a LI-COR 6800 system fitted with 6800-01A fluorometer. All measurements were done under PPFD = 1500 (10% blue vs. 90% red) (µmol m^-2^ s^-1^), chamber temperature at 25°C, ambient chamber CO_2_ concentration (*C*_a_) at 400 (µmol mol^-1^), flow rate at 500 (µmol air s^-1^), and vapor pressure deficit between 1.5-2.0 kPa in five replications per line. All leaves were dark adapted for 20 minutes prior to all other measurements to obtain the maximum quantum yield of photosystem II. The quantum yield of photosystem II (Φ_PSII_) under actinic light was obtained by application of saturating multiphase flashes (>8000 µmol m^-2^ s^-1^) as per Genty *et al*. (1989).

##### 2.2.2. Mesophyll conductance (*g*_m_)

The “variable *J* method” was used to estimate *g*_m_ based on the calculation of electron transport rate (*J*_flu_) from measurements of chlorophyll fluorescence (Bongi & Loreto, 1989; Harley *et al*., 1992):

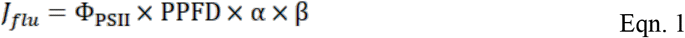

where β (= 0.5 for C_3_ plants) is the fraction of absorbed quanta reaching photosystem II (Bernacchi *et al*., 2002). The leaf absorbance, α, was measured to be 84.2 % based on the average value (± 0.3 standard error) in all individuals using an ASD Fieldspec spectroradiometer (ViewSpec Pro, ASD Inc. Boulder, CO, USA). Mesophyll conductance was given by (Harley *et al*., 1992):

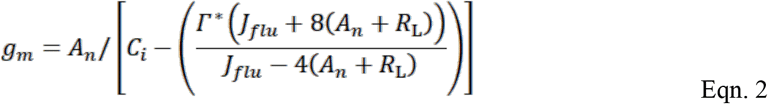

where *R*_d_ is the non-photorespiratory respiration rate in the light, and *Γ\** is the chloroplast CO_2_ photocompensation point. *Γ\** was assumed to equal the intercellular CO_2_ photocompensation point (*C*_i_*) per Gilbert *et al*. (2012). *R*_d_ (0.41 ± 0.05 µmol m^-2^ s^-1^) and *C*_i_* (44.66 ± 0.32 µmol mol^-1^) were estimated using the Laisk method (Laisk, 1977 in Gilbert *et al*., 2012) as the point of intersection of the linear portion of averaged four sets of *A*_n_-*C*_i_ curves obtained at three irradiances (100, 200 and 500 µmol m^-2^ s^-1^) and 13 CO_2_ concentrations (35, 40, 50, 60, 70, 80, 90, 100, 110, 120, 140, 160, and 180 µmol mol^-1^). Having obtained *g*_m_ by the chlorophyll fluorescence method, the CO_2_ concentration in the chloroplast (*C*_c_) was estimated according to Harley *et al*. (1992):

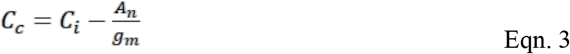

##### 2.2.3. *A*_*n*_*-C*_*i*_ curves

We constructed CO_2_ response (*A*_n_-*C*_i_) curves for lines WT, 9, and 15 under each irrigation treatment at 1500 µmol m^-2^ s^-1^ PPFD under the following sample CO2 concentrations: 40, 50, 80, 100, 150, 200, 400, 500, 600, 700, 800, 900, 1200, 1400 µmol mol^-1^ under well-watered, mild and moderate drought conditions. *A*_n_ and corresponding *C*_i_ values for each curve with 5 replications were introduced to Sharkey’s fitting calculator version 2.0 (Sharkey 2016), a Farquhar-von Caemmerer-Berry (FvCB)-based model, to estimate maximum carboxylation rate (*V*_cmax_) and electron transport rate (*J*) as well as to predict assimilation rates (*A*_n_) at the limiting states of Rubisco (*A*_c_), and RuBP-regeneration (*A*_j_) (Sharkey, Bernacchi, Farquhar & Singsaas, 2007). When triose phosphate use (TPU) limitation was evident, the highest *A*_n_ was used as an estimate of the maximum assimilation rate (*A*_max_) achievable under non-CO_2_-limiting conditions (Sharkey et al. 2007). For validation and enhanced comparison, *g*_m_ was also estimated using *A*_n_-*C*_i_ curves in addition to the independent chlorophyll fluorescence method.

#### 2.3. Leaf hydraulic conductance (*K*_*leaf*_)

Leaf hydraulic conductance was measured following the evaporative flux method (EFM), as previously described for grapevines in Albuquerque *et al*. (2020). Plants were kept well-watered in pots, brought to the lab the day before the measurement, and bagged with wet paper towels to halt transpiration and maintain high humidity. The following morning, shoots were excised and recut under water to prevent air entry into the stems (Wheeler et al., 2013). Shoots with 4-5 mature leaves were left to bench-dry to achieve a range of water potentials. Then each leaf was individually bagged, and placed in a larger bag with wet paper towels for at least 30 min for equilibration. Water potentials were measured using a digital pressure chamber (Model 1000D, PMS Instruments, Albany, Oregon, USA) in the two leaves on each side of the shoot and the averages defined as initial water potential. If the difference was larger than 0.2 MPa, the sample was discarded. For very dehydrated samples, differences of 0.3 MPa were tolerated. One of the remaining leaves was excised under water in a petri dish with ultrapure partially degassed water using a fresh razor blade and inserted in the tubing connected to a water source in the balance. Parafilm was wrapped around the petiole and tubing to secure and prevent any air from getting into the system. The leaf was held above the water level to verify that the sample was secured and placed in the system between two layers of coarse fishing net to hold the leaf in place above the fan and below the light source (500W Halogen Work Light, Ace Hardware, Oak Brook, IL, USA). Photosynthetically Active Radiation (PAR) of 1100 µmol m^-2^ s^-1^ (±200 µmol m^-2^ s^-1^) was measured with a quantum sensor (190R, LiCor Biosciences, Lincoln, NE, USA) with a water bath to avoid overheating as leaves should be between 23°C and 28°C to account for variations in water viscosity. After steady-state was reached with a flow rate <0.05 coefficient of variation of the last ten measurements and of the last five measurements for very low flow rates and no trend upwards or downwards, the leaf temperature was measured in the adaxial part with a type T thermocouple and immediately taken from the system and placed in plastic bag which had been previously exhaled in to saturate with CO2 and humidity. Then this bag was placed in the dark in a larger bag with wet paper towel and final leaf water potential was measured after at least 30 min of equilibration. Leaf area was measured with a leaf area meter (LI-3000C, LiCor Biosciences, Lincoln, NE, USA) and used to normalize *K*_leaf_ to the last five minutes of steady-state flow-rate measurements, with water viscosity calculated at 25 °C (Scoffoni et al., 2012).

### 3. Statistical analysis

#### 3.1. Aquaporin expression

Relative expression data were analyzed with three-way ANOVA, with Tukey’s multiple comparisons test (α = 0.01) to evaluate the effects of irrigation (well-watered, mild and moderate drought), genotype (WT, 9, 15) and tissue type (petioles and leaves). Due to the nature of the relative expression data, variables were transformed by the natural logarithm to meet ANOVA’s assumptions. Statistical analyses were performed using the InfoStat software program, version 2.0 (Di Rienzo et al. 2014). Only significant results are shown.

#### 3.2. Leaf gas exchange

A two-way ANOVA with Tukey’s multiple comparisons test (α = 0.01) was used to analyze irrigation and genotype effects for *A*_n_, *g*_s_, and *g*_m_ using instantaneous gas exchange measurements, Ψ_leaf-midday_, Ψ_leaf-predawn_ and *A*_max_, *V*_cmax_ and *J* derived from the Sharkey calculator v2 using *A*_n_-*C*_i_ curves (GraphPad Software, Inc. CA, USA).

#### 3.3. Leaf hydraulic conductance

To find the best fit model for each *K*_*leaf*_ vulnerability curve, a protocol was followed as previously described in Scoffoni et al. (2012). A Dixon’s test (α < 0.05) was performed in a dataset in line to detect any outliers before each of four functions were tested as the best fit model using R Studio. We tested a linear function (Eqn. 4); a three-parameter sigmoidal function (Eqn. 5); a three-parameter logistic function (Eqn. 6) and a three-parameter exponential function (Eqn. 7):

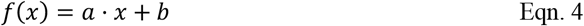

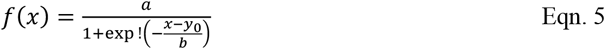

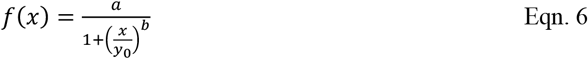

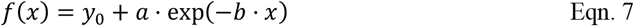

The function with the lowest Akaike Information Criterion (AIC) corrected for low *n* was chosen, unless the difference was <2, when the function with the highest *r*^2^ was the best fit model. We compared line responses by fitting nested nonlinear models to the combined dataset (common curve vs. line-specific vs. fully line-specific parameters) and compared them by analysis of variance on residual sums of squares. Parameter uncertainty and 95% confidence intervals (CIs) for maximum *K*_leaf_ and 50% and 80% loss with dehydration (*K*_max_, P_50_*K*_leaf and_ P_80_*K*_leaf_, respectively), and their pairwise differences among lines were estimated by parametric bootstrap. Analysis was carried out using R Studio and plots were generated with package ggplot2.

## RESULTS

### Gene expression

Both mutants exhibited similar expression of *PIP2;1* and significantly lower expression than the WT under all conditions (Fig. 2). Leaf lamina showed greater PIP2;1 expression at predawn (P=0.002) and midday (∼5-fold; P<0.0001) than petioles under well-watered conditions. Under drought at midday, PIP2;1 expression in the lamina decreased significantly (P<0.0001) and was lowest for mutants. In petioles, no differences in *PIP2;1* expression were observed between the water treatments among the two mutants and WT (Fig.2). Mutants exhibited low *PIP2;1* expression at both predawn and midday, while WT maintained the same expression at pre-dawn between watering treatments. We did not observe overexpression of other aquaporins (e.g *PIP1;1*; *PIP2;2*; *PIP2;3, TIP1;1*; and *TIP2;1*) to compensate for the knock-out of *PIP 2;1* (Sup. Fig. S1).

**Figure 2.**
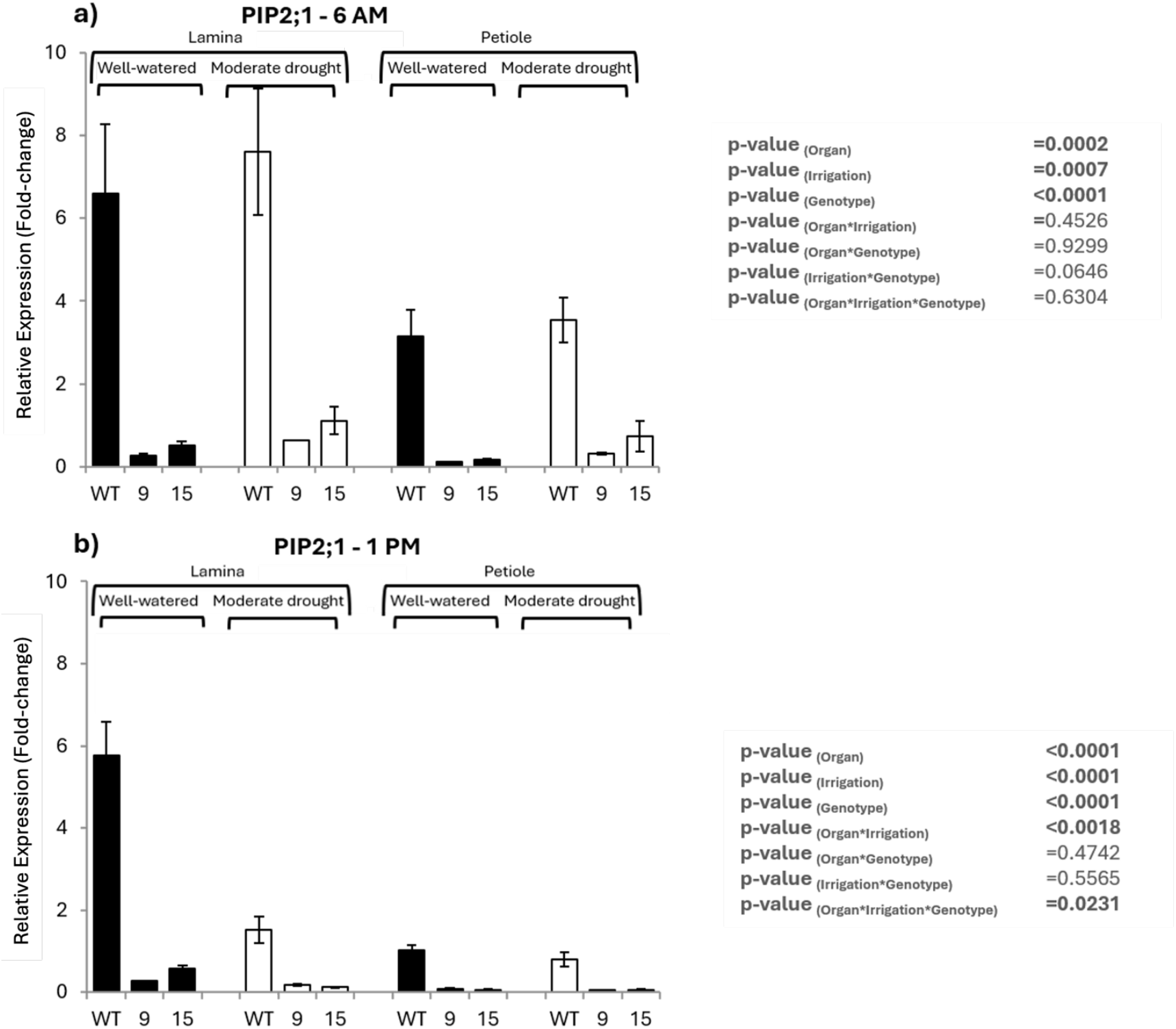
Relative expression of *PIP2;1* (2^(-△△Ct)) measured by qPCR, in petioles and leaf laminas of well-watered and moderate drought in Chardonnay mutant lines 9 and 15 and wild type (WT). Relative expression levels are shown at 6am for pre-dawn (a) and 1pm for midday (b). Fold changes were measured relative to the levels detected in well water WT petiole at 1 PM. Bars represent means and standard errors. ANOVA p-values are shown on the right.

### CO_2_ Response Curves

Under well-watered and mild drought conditions, all lines exhibited similar maximum carboxylation rates (*V*_cmax_) and maximum net assimilation rates (*A*_max_). However, under moderate drought, lines 9 and 15 showed significantly higher *V*_cmax_ and *A*_max_ compared to the WT (Fig. 3, Fig. S2).

**Figure 3.**
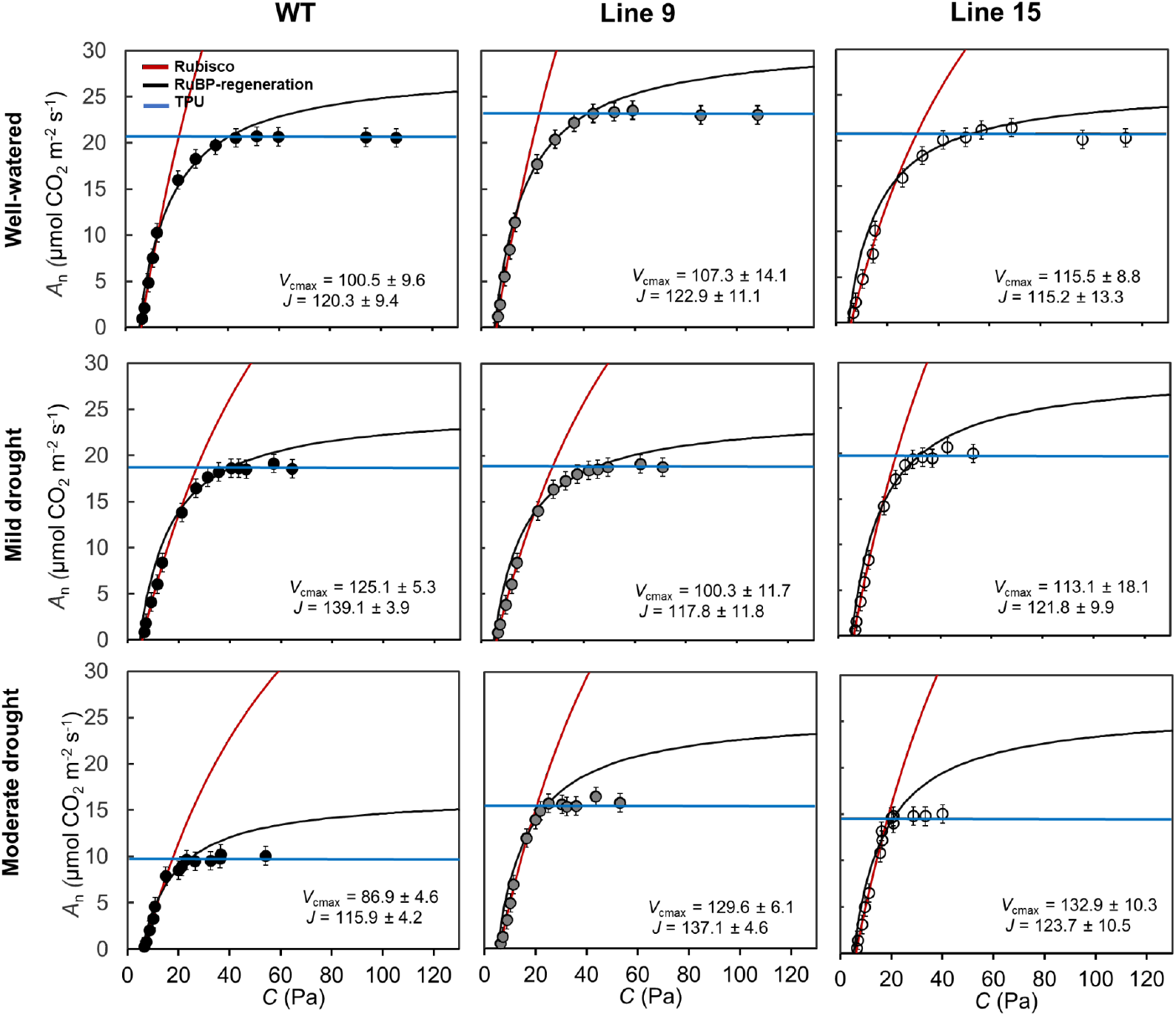
Photosynthetic CO_2_ response curves were constructed using Sharkey’s fitting calculator version 2.0 (Sharkey 2016), averaged over five replications for WT (black circles), line 9 (gray circles) and line 15 (empty circles) under well-watered, mild and moderate drought conditions and error bars (± SE, n = 5) were measured directly. *A*-*C*_i_ curves were used to generate *V*_cmax_ and *J* and averaged over five replications for each line under different treatments (± SE, n = 5). Blue horizontal lines indicate assimilation rate at triose phosphate use limitation matching the assimilation rate at the CO_2_ saturating state (*A*_max_), while Rubisco and RuBP regeneration limitations are indicated for each line by red and black curves, respectively.

### Gas exchange under ambient CO_2_

Net assimilation rate (*A*_n_) and stomatal conductance (*g*_s_) were comparable across all lines under well-watered and mild drought conditions. However, under moderate drought, *A*_n_ and *g*_s_ were significantly lower in the WT than lines 9 and 15 (*P* < 0.001; Fig. 4). In contrast, mesophyll conductance (*g*_m_), showed no significant differences between lines under any condition (Fig. 4). The *g*_m_ estimates from the two different methods, chlorophyll fluorescence and *A*_n_-*C*_i_ curves, were significantly correlated (*r*^2^ = 0.61, *P* = 0.01) (Fig. S3). Consistent with the findings for *g*_m_, intrinsic water use efficiency (WUE_i_) also did not differ significantly between the mutant lines and WT under either mild or moderate drought conditions (Fig. S4).

**Figures 4.**
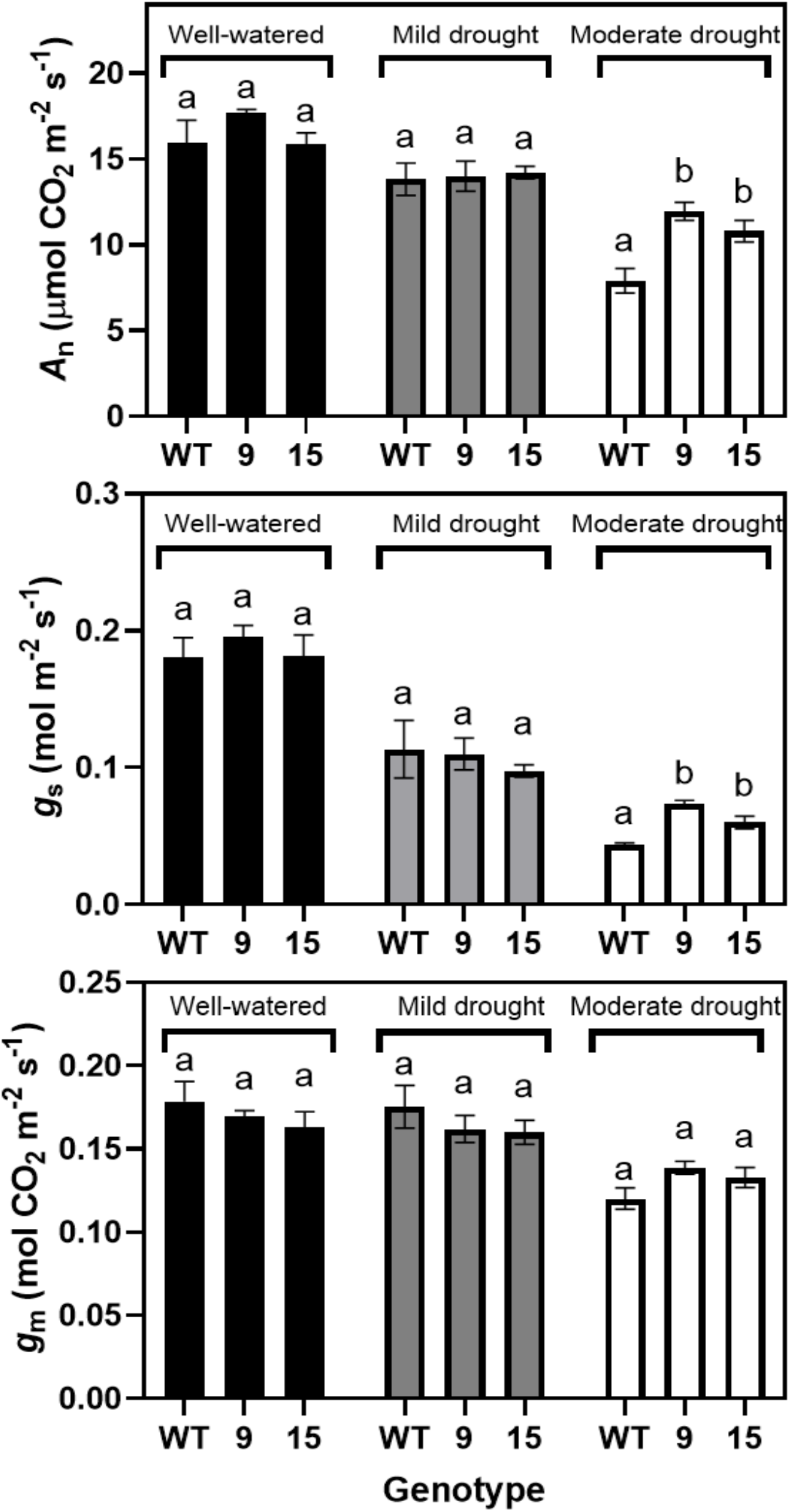
Net assimilation rate (*A*_n_, μmol CO_2_ m^-2^ s^-1^), stomatal conductance (*g*_s_, mol m^-2^ s^-1^), and mesophyll conductance (*g*_m_, mol CO_2_ m^-2^ s^-1^) for WT (black), line 9 (gray) and line 15 (white) under well-watered, mild drought and moderate drought conditions. The measurements were averaged over five replicates per line and irrigation treatment combination (± SE, n = 5).

### Leaf Water Potential

Midday leaf water potential (Ψ_leaf-midday_) differed significantly between the WT and lines 9 and 15 only under moderate drought conditions (Fig. 5). Under this stress, the mutant lines exhibited more negative Ψ_leaf-midday_ values than the WT, which reached -1.2 MPa. In contrast, predawn leaf water potential (Ψ_leaf-predawn_) showed no significant differences between the WT and mutant lines across all treatments.

**Figures 5.**
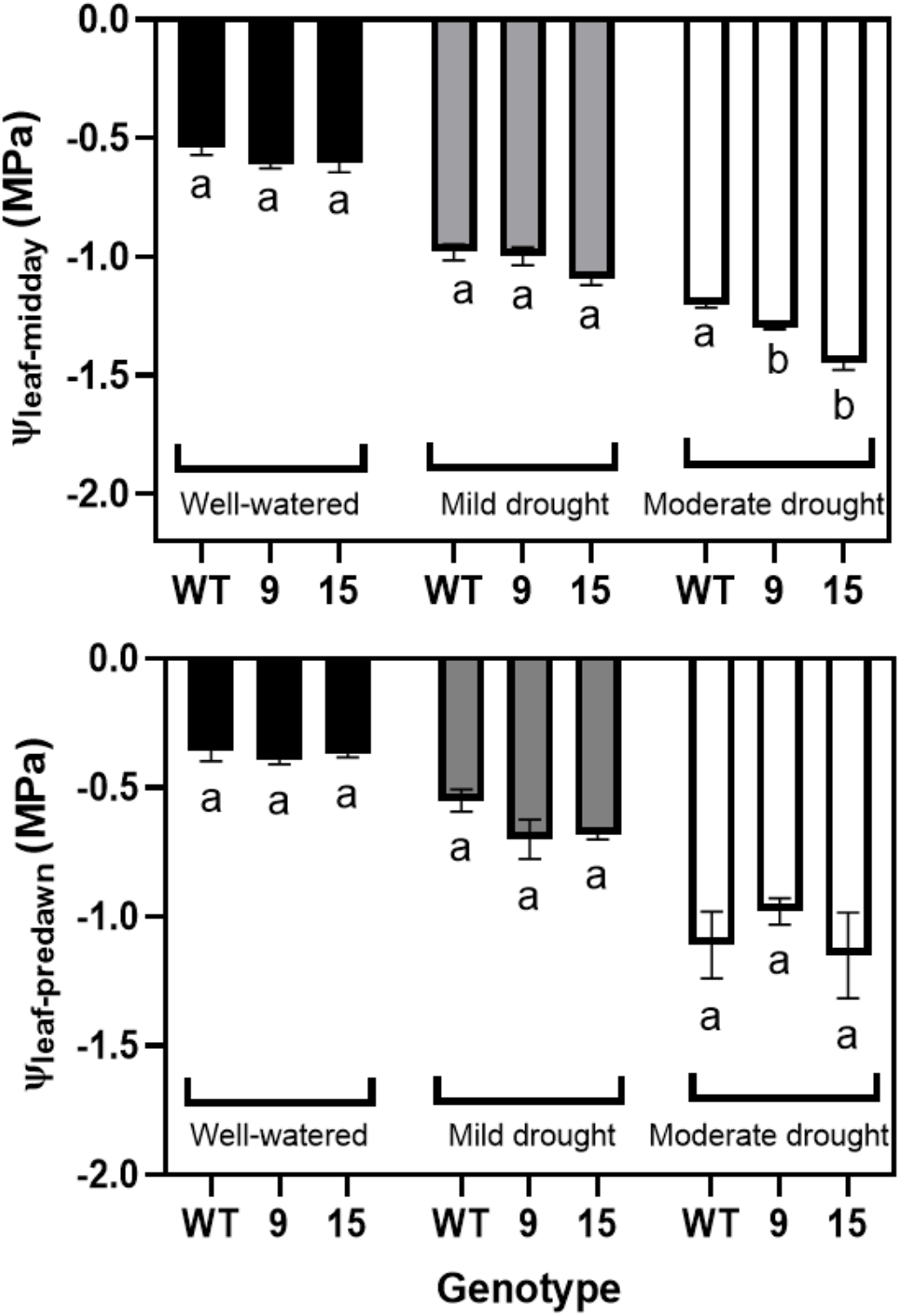
Midday and pre-dawn leaf water potentials (Ψ_leaf-midday_ and Ψ_leaf-predawn_) for WT (black), line 9 (gray) and line 15 (white) under well-watered, mild drought and moderate drought conditions. The measurements were averaged over five replicates per line and irrigation treatment combination (± SE, n = 5).

### Leaf Hydraulic Conductance (K_leaf_)

All lines exhibited similar *K*_leaf_ vulnerability with dehydration with exponential responses (R^2^ = 0.95, 0.92 and 0.88 for WT, lines 9 and 15, respectively; all P < 0.0001 (Fig. 6; Table 2). The water potential at 50% loss of *K*_*leaf*_ (P_50_*K*_*leaf*_) ranged from Ψ_w_ = -0.33, -0.34 and to - 0.25MPa, and at 80% loss of *K*_*leaf*_ (P_80_*K*_*leaf*_) from Ψ_w_ = -0.75, -0.76 to -0.60MPa for the WT, lines 9 and 15, respectively. In the same order, *K*_*max*_ was 16.0, 18.6 and 16.7 mmolH_2_O m^-2^ s^- 1^MPa^-1^ for the WT, line 9 and 15. Global nonlinear model comparison indicated differences in *K*_*max*_ and vulnerabilities across lines (P = 0.009 and P = 0.046), however, bootstrap 95% CIs for pairwise differences in *K*_*max*,_ P_50_*K*_*leaf*_, P_80_*K*_*leaf*_ spanned zero. We thus interpreted any differences in *K*_*leaf*_ vulnerability as small and consider the three lines to have similar responses.

**Table 2.** Leaf hydraulic conductance (*K*_leaf_) for WT and lines 9 and 15 of *V. vinifera*. Values of maximum *K*_leaf_ and vulnerability curves showing water potentials at which *K*_leaf_ declined by 50% (P_50_*K*_leaf_) and 80% (P_80_*K*_leaf_). Values are model estimates with 95% confidence intervals (CIs) from parametric bootstrap (n = 2000). CIs overlapped among lines, indicating only small quantitative differences.

| *V. vinifera* cv.<br>Chardonnay | $K_{\text{leaf}}$ | | |
| --- | --- | --- | --- |
| | $K_{\text{leaf}}$ maximum<br>(mmolH<sub>2</sub>O m<sup>-2</sup> s<sup>-1</sup>MPa<sup>-1</sup>) | $P_{50}K_{\text{leaf}}$<br>(MPa) | $P_{80}K_{\text{leaf}}$<br>(MPa) |
| WT | 16.0 (10.8–21.3) | −0.33 (−0.53 to −0.23) | −0.75 (−1.11 to −0.56) |
| Line 9 | 18.6 (13.1–24.1) | −0.34 (−0.50 to −0.24) | −0.76 (−1.06 to −0.59) |
| Line 15 | 16.7 (14.0–21.9) | −0.25 (−0.33 to −0.20) | −0.60 (−0.76 to −0.49) |

**Figure 6.**
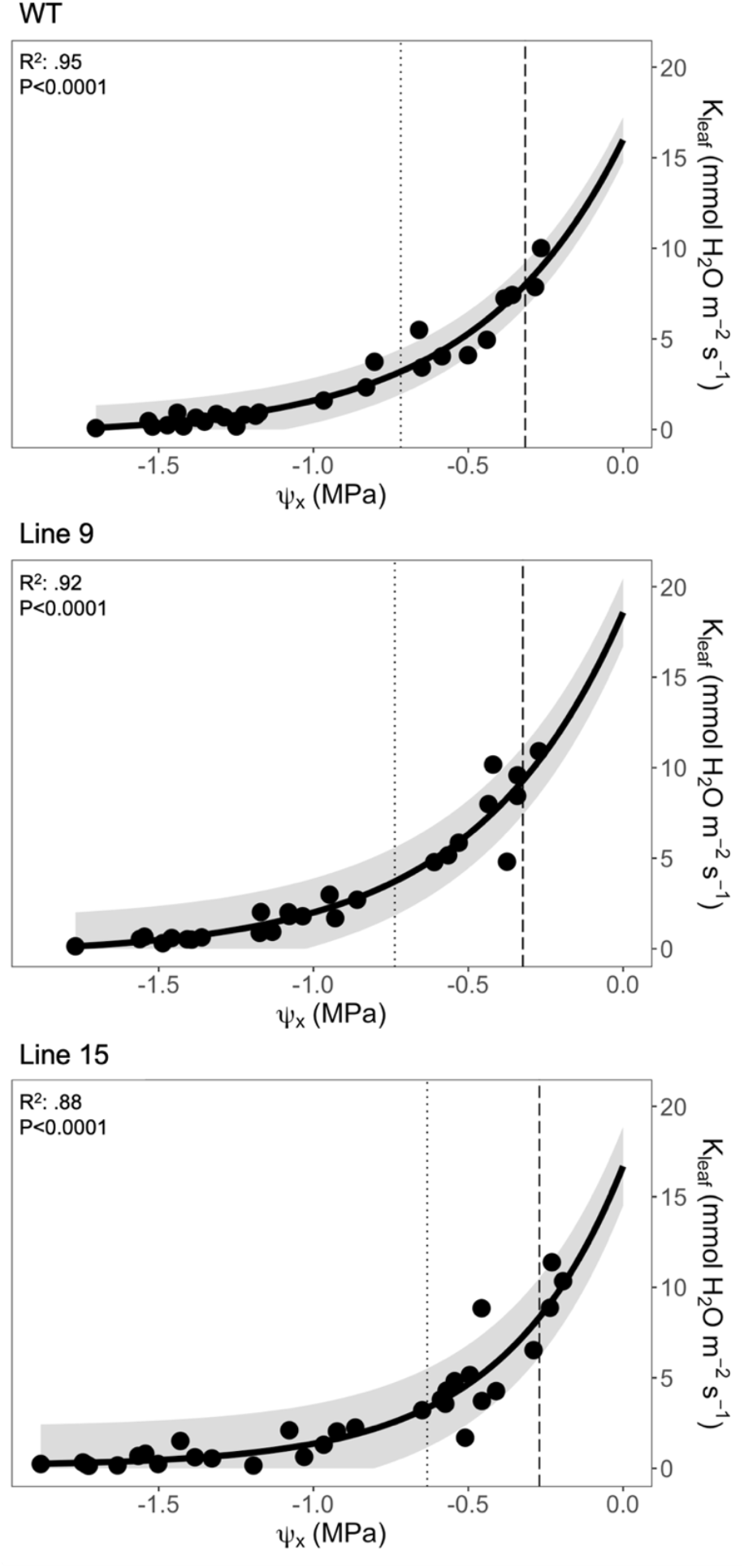
Leaf hydraulic conductance (*K*_leaf_) vulnerability curve of the different Chardonnay of WT and mutant lines. Points show measured data and lines as the best fit curve using a three-parameter exponential function. Shaded area indicates 95% confidence interval (CI) from parametric bootstrap. Vertical dashed and doted lines indicate P_50_*K*_leaf and_ P_80_*K*_leaf,_ respectively. Curve shapes and position indicate similar *K*_leaf_ responses despite statistical differences detected in statistical models (Table 2 and Materials and Methods). Best fitted curves were three-parameter exponential: y=-0.3418+ 16.32953e–(– 2.1296)Ψ_w_, n=26 (WT); y= -0.338+ 18.9203e–(–2.0886)Ψ_w_, n=26 (b); y= 0.1274 + 16.5657e–(– 2.5931)Ψ_w_, n=31 (c). Gray lines around the fitted curve are the confidence intervals.

## DISCUSSION

Our findings provide evidence that *PIP2;1* plays an important role in regulating grapevine responses to water stress, primarily by inducing stomatal closure which, in turn, reduces photosynthetic carbon capture. By combining physiological measurements of photosynthesis and leaf hydraulics during dehydration with novel molecular techniques of gene editing like CRISPR-Cas9, our work provides unexpected insights into how an aquaporin isoform can regulate leaf function in grapevine under water stress. We chose to target *PIP2;1* as previous work identified this isoform as playing an important role in grapevine responses to water stress (Gambetta et al.; 2013; Pou et al. 2013; Chitarra et al. 2014; Lukšić et al. 2023; Braidotti et al. 2024).

Mutations were produced using the CRISPR-Cas9 editing tool and were confirmed through sequencing. Because the two targeted sites exhibited differences in the mutations produced in lines 9 and 15 (Table 1), it is expected that the magnitude of the editing effect on the *PIP 2;1* gene transcription/translation and/or protein function might also differ. The expression of *PIP2;1* varied with organ, time and water status in WT, with the highest expression in the lamina, that strongly decreased at midday under water stress conditions. Mutants showed little to no *PIP2;1* expression under all conditions. Like our midday results, another study observed down-regulation of *VvPIP2;1* expression in petioles of Chardonnay in response to water stress, with reduced expression in leaf laminas of both Chardonnay and Grenache with more negative Ψ_leaf_ (Shelden et al., 2017).

A remarkable role of *PIP2;1* mutation was found on stomatal conductance. The greater photosynthetic capacity of lines 9 and 15 was accompanied with an increased stomatal conductance under moderate drought conditions of this experiment in comparison to the wild type. However, there was no significant difference in mesophyll conductance between lines under any condition. Even though *V*_cmax_ and *A*_n_ did not differ among lines under well-watered and mild drought, under moderate drought, both were higher in the mutants (Fig. 3). This suggests that the WT exhibited stronger down-regulation of photosynthetic capacity (biochemical limitation) rather than stomatal limitation than the mutants under moderate drought. Indeed, drought can progressively down-regulate photosynthetic metabolism, so targeted measurements of Rubisco activation/state and electron transport capacity would help identify the mechanisms underlying the *V*_cmax_ differences (Flexas and Medrano, 2002). Aquaporins have also been shown to facilitate CO_2_ diffusion across membranes and influence photosynthesis in some systems (Heckwolf et al., 2011; Uehlein et al., 2012); however, the similar *g*_m_ responses across our lines do not support a dominant role of *PIP2;1* in mesophyll CO_2_ diffusion in our study.

This points to *PIP2;1* playing a pivotal role in stomatal closure during drought. Arabidopsis *AtPIP2;1* has been shown to be essential for stomatal closure in response to the drought-induced hormone abscisic acid (ABA) (Grondin et al. 2015). ABA-triggered stomatal closure requires an increase in guard cell permeability to water and possibly hydrogen peroxide, through the OST1-dependent phosphorylation of *PIP2;1* at Ser-121. *PIP2;1* is also implicated in the influx of apoplastic H_2_O_2_ to the cytosol during ABA-dependent stomatal closure (Grondin et al. 2015). Overexpression of *VvPIP2;4N* in grapevine roots increased stomatal conductance in well-watered grapevines, however, higher ABA accumulation under drought counteracted this effect and lead to similar stomatal closure as in wild-type plants (Perrone et al., 2012). We observed that leaf laminas of the WT maintained high expression of *PIP2;1* at pre-dawn and midday under well-watered conditions, indicating that factors like turgor and ABA synthesis could contribute to the down-regulation of aquaporins and stomatal conductance.

In another recent study, Arabidopsis *AtPIP2;1* was shown to facilitate H_2_O_2_ entry into guard cells, leading to stomatal closure in response to ABA or the bacterial elicitor flg22. Rodrigues et al., (2017) showed that accumulation of H_2_O_2_ into guard cells in response to either flg22 and ABA was abolished in site-directed mutants of Arabidopsis *PIP2;1*. They also showed that the intrinsic transport activity of *AtPIP2;1* strongly relies on *AtPIP2;1* phosphorylation by the guard cell SnRK2;6 (OST1) protein kinase. A phosphorylation-dependent switch between ion and water permeation in *AtPIP2;1* might be explained by coupling between the gates of the four monomer water channels and the central pore of the tetramer (Tyerman et al. 2021). The involvement of *PIP*s in stomatal dynamics has been documented in several studies showing their role in facilitating the transport of H_2_O, H_2_O_2_ and CO_2_ during stomatal closure induced by ABA and CO_2_ in sunflower, maize, tomato and poplar (Ding and Chaumont 2020, Ding et al, 2024). The tighter control of stomatal conductance of the WT at moderate drought (Fig. 4) can explain the less negative water potentials in comparison with the mutants (Fig. 5). This earlier stomatal closure driven by *PIP2;1* prevented xylem tension in the WT to increase to the brink of water potentials that can lead to inception and rapid spread of vein xylem embolisms in grapevines including Chardonnay (Albuquerque et al., 2020). On the other hand, overexpression of a tonoplast aquaporin in tomato promoted higher transpiration and yield under water and salt stress (Sade et al., 2009). These contrasting results indicate that manipulating different aquaporin isoforms and membrane compartments can shift the balance between hydraulic safety and water use for different crops and environmental conditions, highlighting aquaporins as promising tools to improve crop responses and resilience to drought and heat events when water availability is limited.

Contrary to the differences observed in stomatal responses, WT and knockouts showed similar *K*_leaf_ vulnerabilities to dehydration (Table 2 and Fig. 6). These results were surprising since aquaporins are pointed as the regulators of leaf water transport in the outside-xylem pathways before xylem embolism (Scoffoni et al. 2017; 2018; Albuquerque et al. 2020). Indeed, aquaporin deactivation has been shown to drive *K*_leaf_ decline during drought in several species (Kim and Steudle, 2007; Shatil-Cohen *et al*., 2011; Pantin *et al*., 2013; Pou *et al*., 2013; Sade *et al*., 2014, 2015). Expression of *PIP2;1* was greatly reduced in leaf lamina at midday under moderate drought in the WT (Fig. 2b), with results similar to those previously reported, linking aquaporin expression with vine water status in Chardonnay (Shelden et al. 2017). *K*_leaf_ decline during drought was correlated with reduced *PIP2;1* and *TIP2;1* expression in Chardonnay (Pou et al. 2013). However, the effective knock-down of *PIP2;1* in our mutants and the similar *K*_leaf_ responses to dehydration across all lines suggest that *PIP2;1* was not involved in down-regulating *K*_leaf_ decline during dehydration in grapevines in our study. Martre et al (2002) did not find any differences in *K*_leaf_ between WT and plants with reduced of some *PIP1* and *PIP2* isoforms under well-watered conditions in Arabidopsis.

Our results showed that *PIP2;1* is not related to *K*_leaf_ decline during drought in grapevine leaves, and we found no overexpression of other aquaporin isoforms (Fig. S1) that could have compensated for the function of *PIP2;1* to explain the similar *K*_leaf_ response to drought across the wild-type and the mutants. These findings do not rule out the role of aquaporins in leaf water transport during drought, but rather highlight its complexity and the importance of new technologies such as CRISPR-Cas9 to elucidate the roles of different aquaporin isoforms in plant responses to drought. Although we targeted specifically to knock out *PIP 2;1*, other isoforms could also be involved in responses to water in the leaves and roots of several species Laur & Hacke, 2013; Monneuse et al., 2011; Gambetta et al., 2012; Prado et al., 2013; Sade et al., 2014; Shatil-Cohen et al., 2011). Indeed, aquaporin function can also be buffered by isoform interactions and post-translational regulation, which can alter permeability and selectivity without requiring transcript upregulation (Tyerman et al., 2021). Moreover, tracking the expression of different aquaporin isoforms across the full dehydration range will be important to elucidate their role in driving leaf water transport decline during drought and how this differs across species. Apoplastic pathways were reported to potentially play an increased role in driving *K*_leaf_ under water stress in Chardonnay (Pou et al., 2013). In addition to a possible mixed interpretation of these results as stated by the authors, this scenario would also require the lack of a Casparian-like suberized barrier in the apoplast. Grapevines, however, have been shown to accumulate suberin in response to water stress in roots and petioles (Vandeleur et al. 2009; Hochberg et al. 2013). Future studies integrating localized suberization with targeted expression of aquaporin isoforms will be fundamental to fully elucidate how they regulate leaf water transport across a range of dehydration in different species.

While we did not observe impacts of *PIP2;1* in *K*_leaf_, the role of this aquaporin isoform in regulating stomatal closure during dehydration is clear. The mutants maintained more stomata open as drought progressed, and this lack of control led to higher xylem tension that could induce catastrophic embolism formation. Early stomatal closure in the WT would act as a protective mechanism that prevents further water loss and build-up of risky tensions in the xylem. This aquaporin-mediated response before hydraulic failure contributes to maintain hydraulic safety margins as well as crop drought resilience. Leaves would be able to quickly restore function when conditions improve like lower VPD and/or an irrigation event. Other structural and physiological processes, including ABA synthesis, turgor pressure, xylem collapse and suberization in the bundle-sheath tissues may also contribute to leaf dynamic responses to stress and recovery. Elucidating how these processes interact with aquaporin function will be fundamental to understanding leaf hydraulic function loss and recovery before xylem embolism.

## CONCLUSIONS

Taken together, our results show that gene-editing using CRISPR-Cas9 in combination with physiological measurements is an important tool towards a better understanding of the mechanisms involved in plant responses to drought. The differences observed between mutants and the WT at moderate water stress show that *PIP2;1* aquaporins have a significant effect on regulating stomatal closure and photosynthetic responses, while having a minimal role in regulating *K*_leaf_. Our findings highlight how aquaporins act as a mechanism to maintain leaf water balance by promoting early stomatal closure under water stress, preventing further water loss and high xylem tensions that could lead to catastrophic embolism. Understanding the complexity involved in the dynamics of leaf responses to changing environmental condition is crucial to improve crop water use-efficiency and resilience to extreme weather events. Future research is needed to partition the role of different mechanisms involved in leaf functional dynamics to water stress and recovery.

## AUTHOR CONTRIBUTIONS

CA and MM contributed equally and share first authorship. CA, MM, AJM and CBA designed the study. CBA, MR, and CArc generated and validated CRISPR lines and contributed to data analysis. CA and MM performed physiological measurements and analyses. CA wrote the first draft. All authors contributed to writing and editing the manuscript.

