## Supplementary Data for "*PIP2;1* aquaporin promotes early stomatal closure in grapevine leaves during water stress"

*^3^ Cátedra de Fisiología Vegetal. Facultad de Ciencias Agrarias, Universidad Nacional de Cuyo, Mendoza, M5507, Argentina*

*^4^Department of Biological Sciences, California State University, Stanislaus, One University Circle, Turlock, California, 95382, USA*

*^5^Department of Plant Biology, University of California, Davis, 605 Hutchinson Rd, Davis, CA 95616, USA*

*^6^ Department of Biological Sciences, California State University, Los Angeles, 5151 State University Drive, Los Angeles, California, 90032, USA*

*^7^USDA-Agricultural Research Service, Davis, CA 95616, USA*

^†^Authors contributed equally.

**SUPPLEMENTARY DATA**

**SUPPLEMENTARY FIGURES**

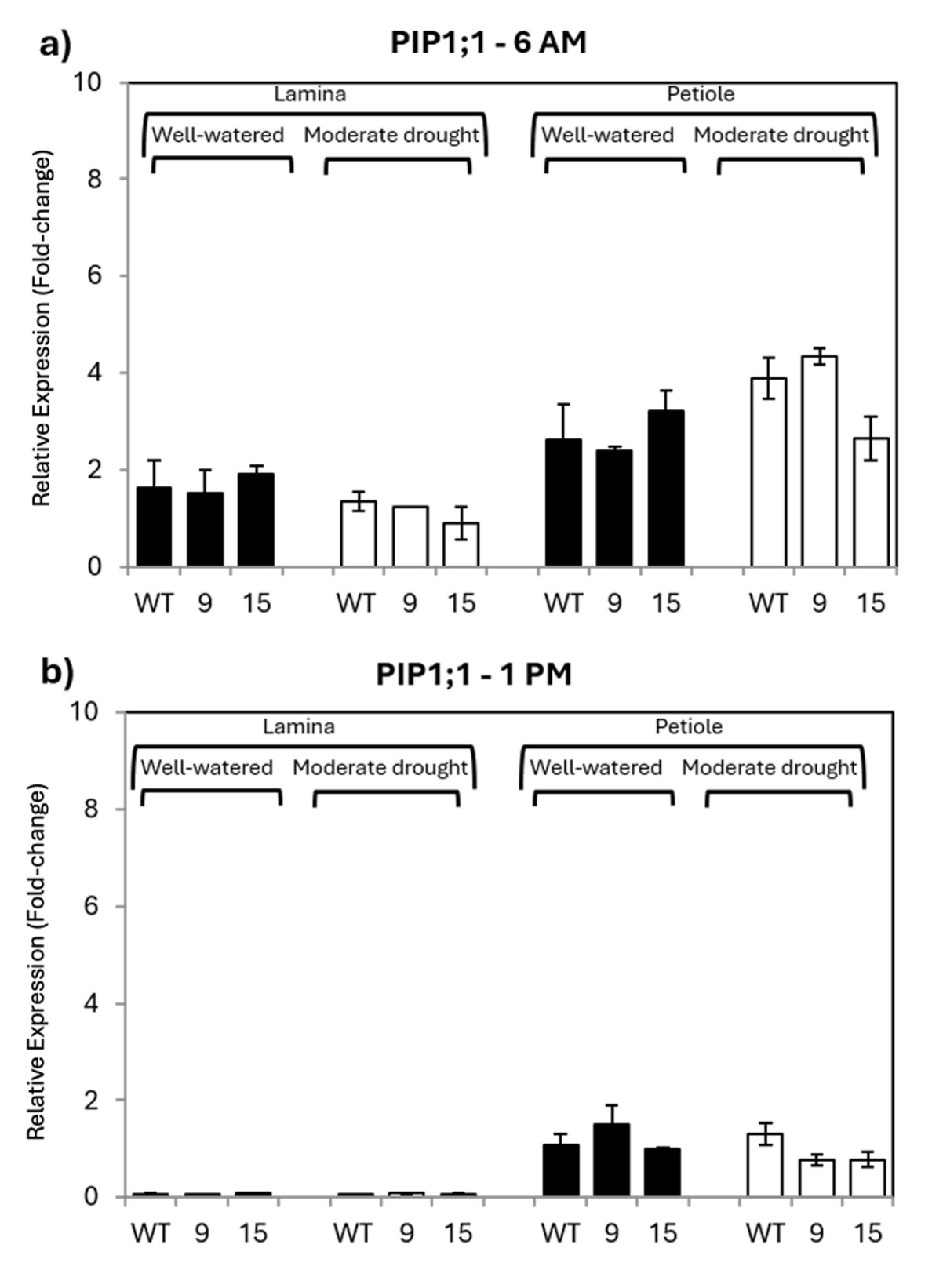

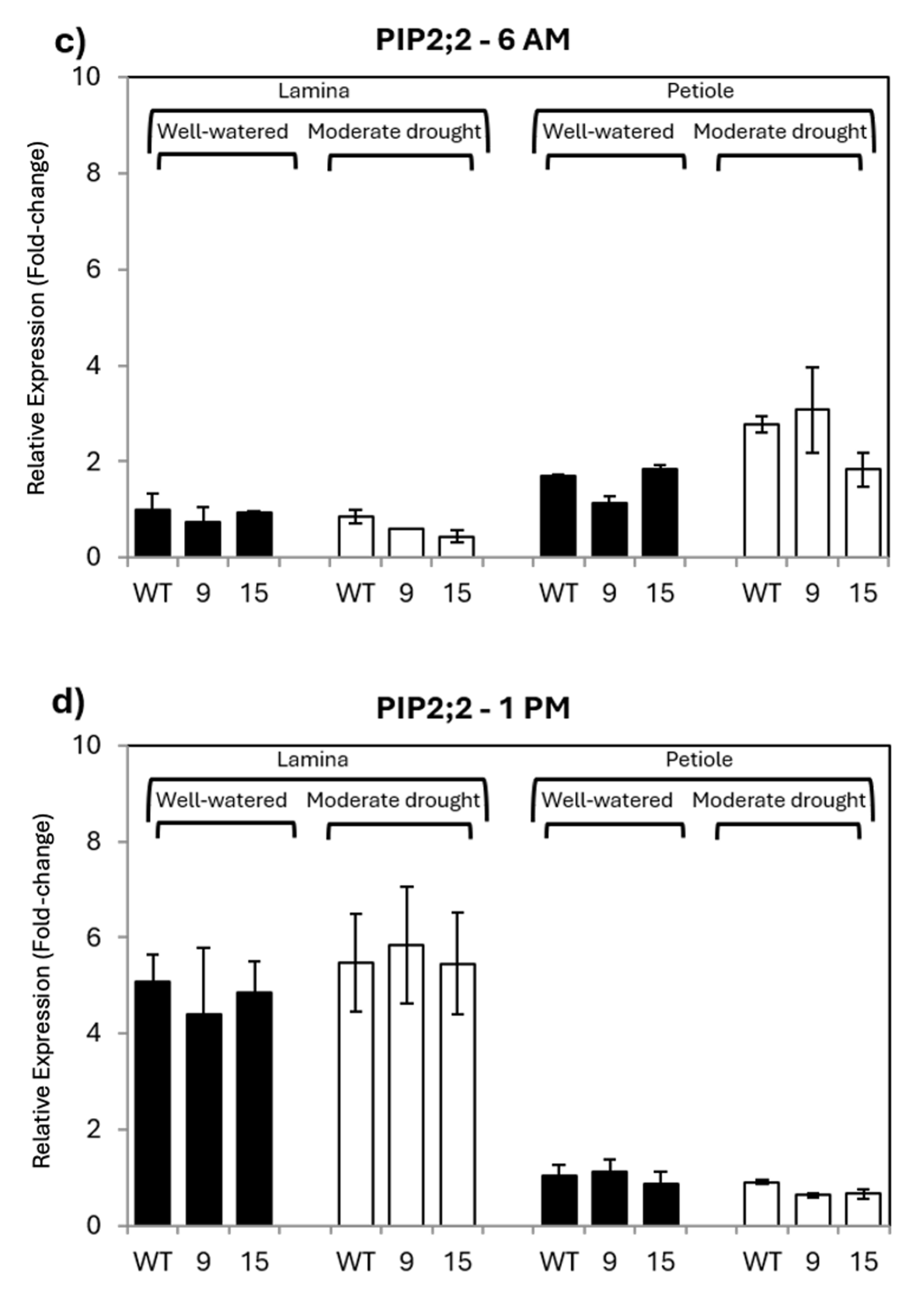

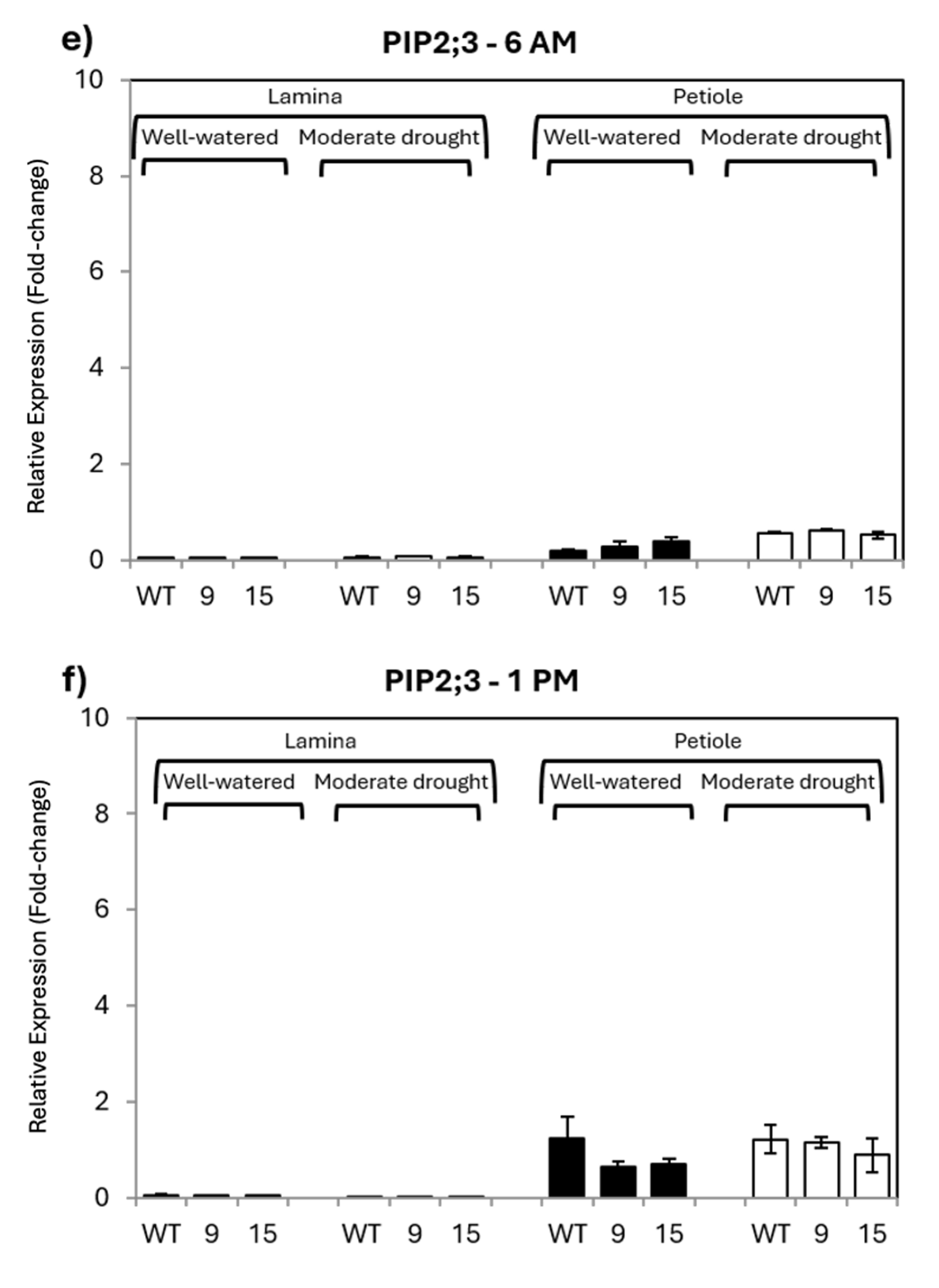

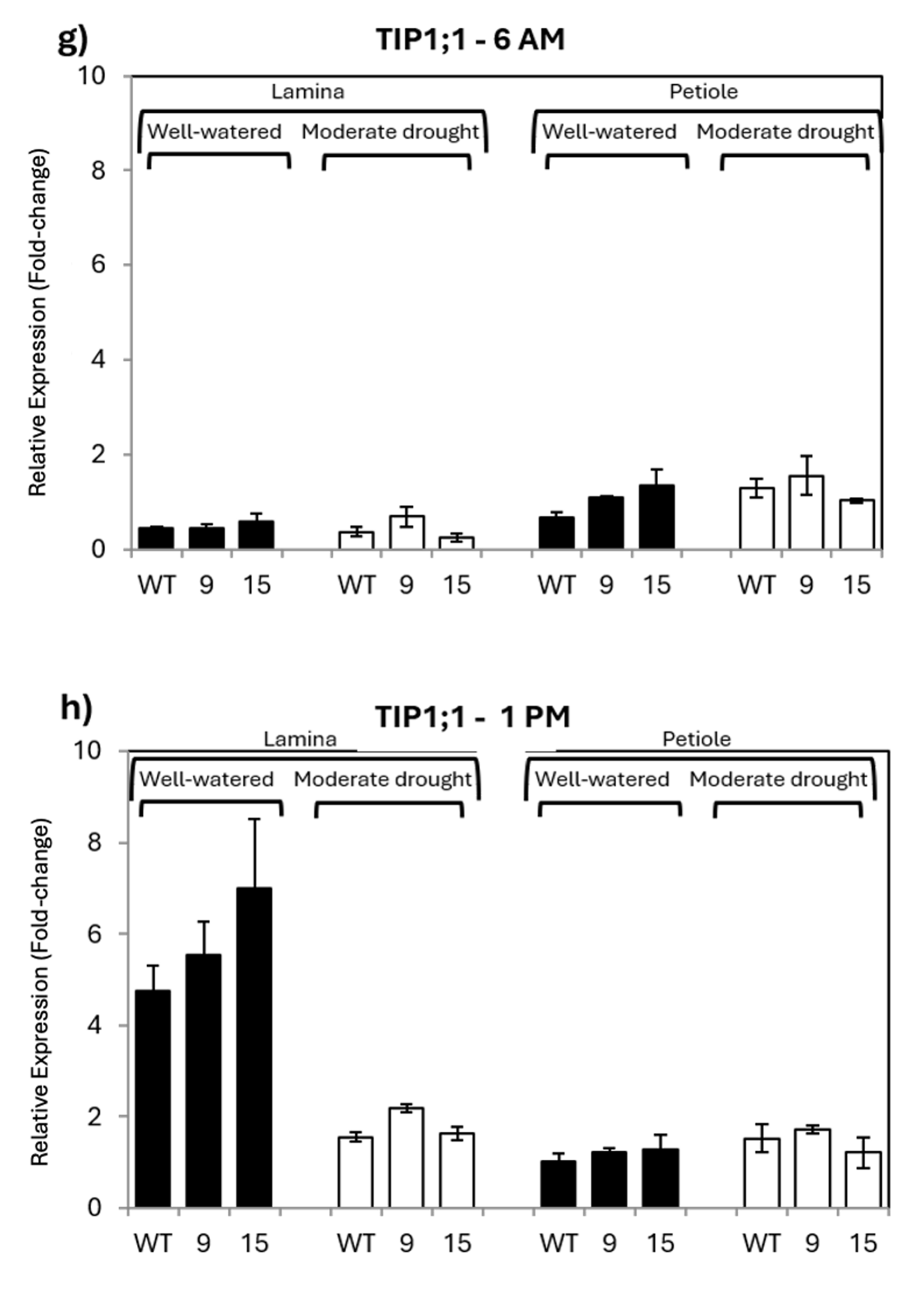

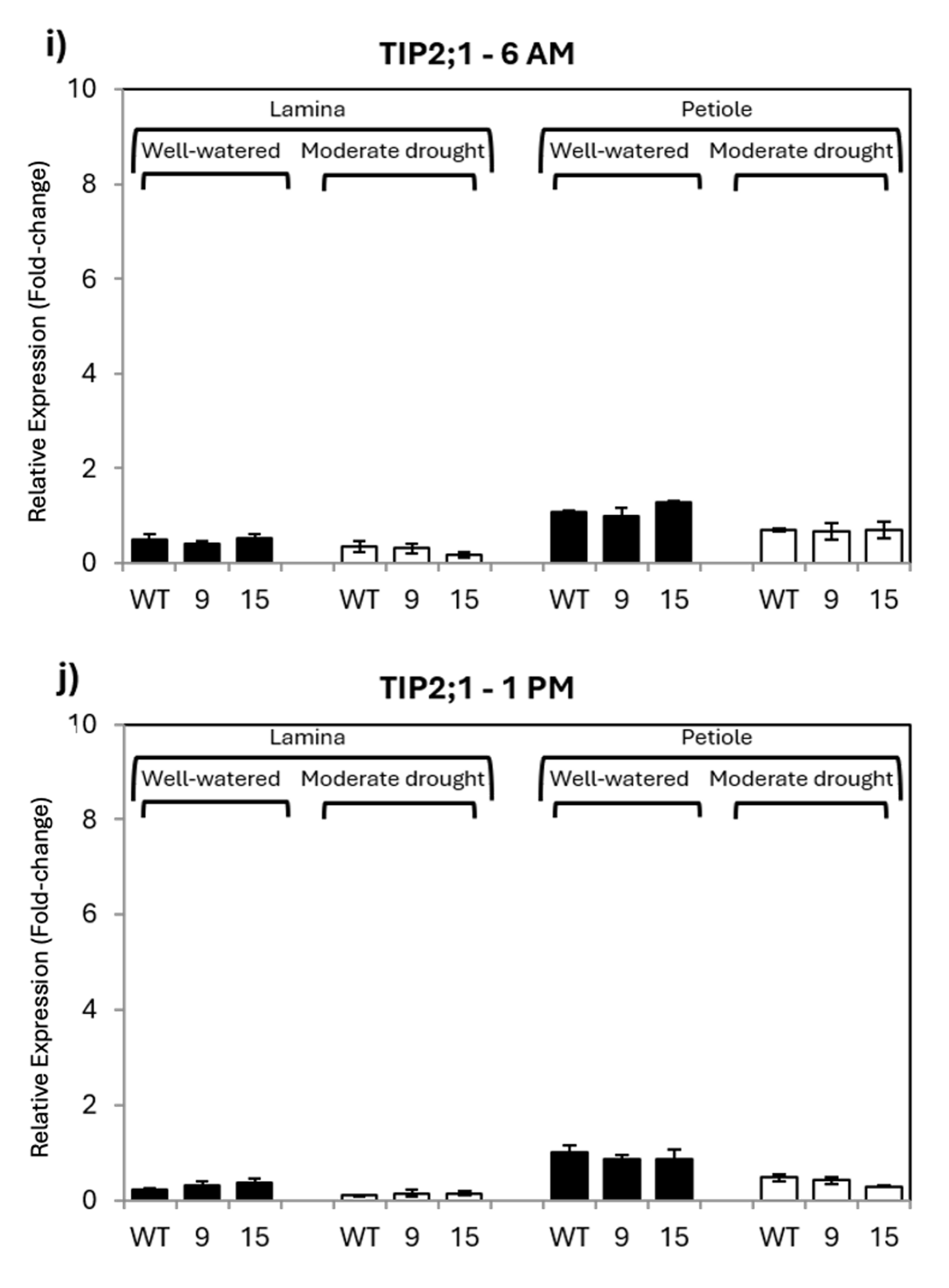

**Figure S1**. Relative expression of *PIP1;1, 2;2, 2;3* and *TIP1;1* and *2;1* (2^(-△△Ct)) measured by qPCR, in well-watered and moderate drought WT and edited lines 9 and 15 of Chardonnay (CH) at 6 AM and 1 PM. Fold changes were measured relative to the levels detected in well-watered WT petiole at 1 PM. Results are reported as means of three biological replicates and three technical replicates. Bars represent standard errors.

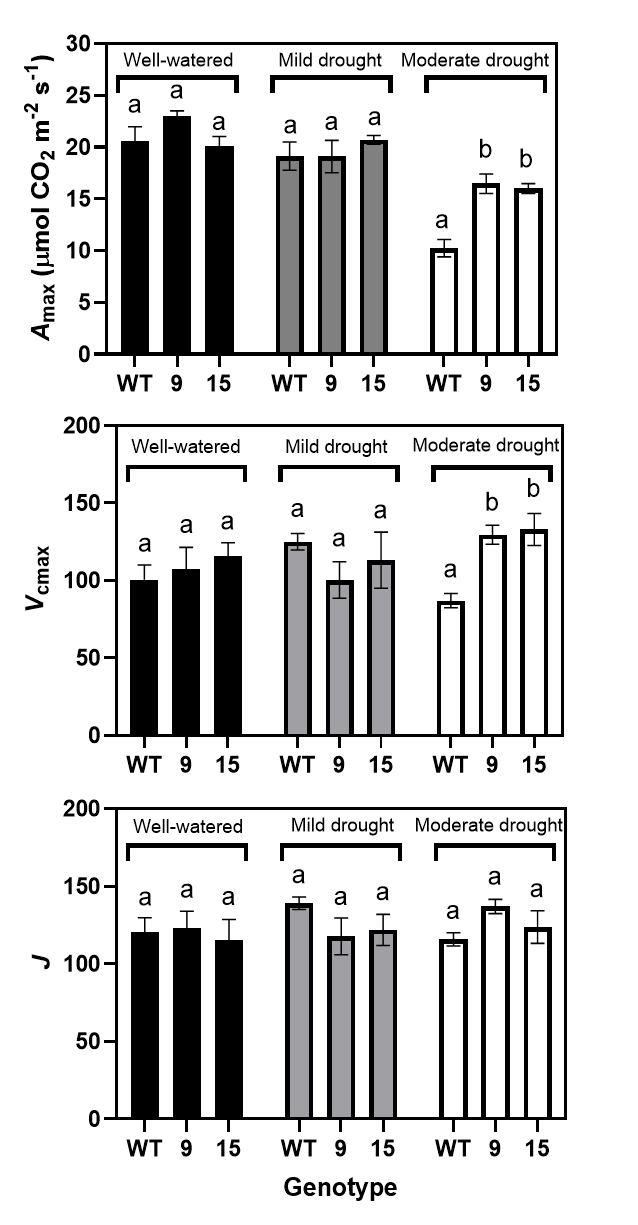

**Figure S2.** Maximum assimilation rate (*A*_max_, μmol CO2 m-2 s-1), Vcmax, and J for WT (black), line 9 (gray) and line 15 (white) under well-watered, mild drought and moderate drought conditions. The measurements were averaged over five replicates per line and irrigation treatment combination (± SE, n = 5). Under moderate drought, mutant lines kept maximum assimilation rate significantly higher than the wildtype line.

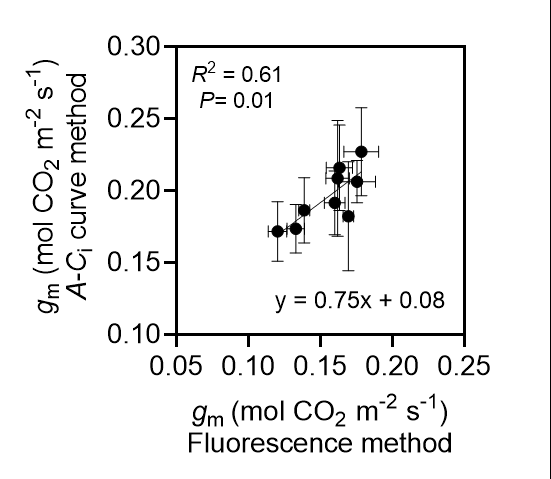

**Figure S3.** Correlation between mesophyll conductance (*g*_m_, mol CO_2_ m^-2^ s^-1^) obtained from chlorophyll fluorescence and *A*_n_-*C*_i_ curve methods for five reps (± SE) for each line under control, mild drought and moderate drought treatments using mean values (± SE, n = 5).

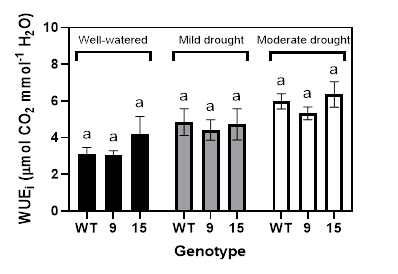

**Figure S4.** Intrinsic water use efficiency (WUE_i_) for WT (black), line 9 (gray) and line 15 (white) under well-watered, mild drought and moderate drought conditions. The measurements were averaged over five replicates per line and irrigation treatment combination (± SE, n = 5). As expected, water use efficiency increased with drought stress. However, there was no significant difference in WUE_i_ between the lines under each water level.
